# An interplay between viruses and bacteria associated with marine sponges from the White Sea revealed by metagenomics

**DOI:** 10.1101/2021.11.12.468366

**Authors:** Anastasiia Rusanova, Victor Fedorchuk, Stepan Toshchakov, Svetlana Dubiley, Dmitry Sutormin

## Abstract

Sponges are remarkable holobionts harboring extremely diverse microbial and viral communities. However, the interactions between the components within holobionts and between a holobiont and environment are largely unknown, especially for polar organisms. To investigate possible interactions within the sponge-associated communities and between them, we probed the microbiomes and viromes of cold-water sympatric sponges *Isodictya palmata, Halichondria panicea*, and *Halichondria sitiens* by 16S and shotgun metagenomics. We showed that the bacterial and viral communities associated with these White Sea sponges are species-specific and different from the surrounding water. Extensive mining of bacterial antiphage defense systems in the metagenomes revealed a variety of defense mechanisms. The abundance of defense systems was comparable in the metagenomes of the sponges and the surrounding water, thus distinguishing the White Sea sponges from those inhabiting the tropical seas. We developed a network-based approach for the combined analysis of CRISPR-spacers and protospacers. Using this approach, we showed that the virus-host interactions within the sponge-associated community are typically more abundant than the inter-community interactions. Additionally, we detected the occurrence of viral exchanges between the communities. Our work provides the first insight into the metagenomics of the three cold-water sponge species from the White Sea and paves the way for a comprehensive analysis of the interactions between microbial communities and associated viruses.

## 1. Introduction

From the nowadays scientific prospect, the majority of multicellular organisms, such as plants and animals, are considered as a combination of the host and the associated communities of microbes and viruses rather than individual units. Together, they make up “holobionts” [1] or “metaorganisms” [2]. Members of the microbial community contribute to the nutrition, protection, immunity, and development of the host they inhabit [3].

One of the most ancient and remarkable examples of holobionts are sponges (Porifera) that can host diverse communities of bacteria, archaea, microalgae, unicellular fungi, as well as viruses and virus-like particles. Associated organisms can occupy up to 40 % of the sponge volume [4], inhabit their mesohyl matrix [3], or localize intracellularly [5]. The composition of sponge-associated microbial communities depends on the host phylogeny rather than on its geographical location and differs from surrounding seawater [6][7][8]. The most frequently detected bacterial phyla in sponges are Proteobacteria (mainly Alpha-, Delta-, and Gamma-), Acidobacteria, Actinobacteria, Bacteroidetes, Chloroflexi, Cyanobacteria, Nitrospirae, Thaumarchaeota [9–11], along with *Candidatus* phylum Poribacteria, which was first discovered in sponges [12].

The majority of sponges are marine, they lead an attached lifestyle and typically feed on microscopic particles by filtering them from surrounding waters [13]. Since a sponge can pump thousands of liters a day, it can absorb a huge number of viruses thus causing a potential threat for associated microbial communities [14][15]. This attracts attention to the metagenomics of sponges, especially for the organisms from tropical and temperate regions. At the same time, sponges from polar regions remained understudied [10]. An expanding corpus of data indicates that genomes of sponge-associated bacteria are enriched with genes of diverse defense systems providing resistance against mobile genetic elements. Adaptive immune systems comprised of clustered regularly interspaced repeats (CRISPR) and CRISPR-associated proteins (Cas proteins), restriction-modification systems (RM), toxin-antitoxin systems (TA), DNA phosphorothioation system (DND), phage exclusion systems (Pgl), DISARM, and Zorya were found to be enriched in comparison to the metagenomes of the surrounding seawater [16–21].

Viroms of several sponges from temperate and tropical locations were investigated [15,22–26]. In all studies, the most commonly identified viruses were tailed dsDNA bacteriophages belonging to Podoviridae, Siphoviridae, and Myoviridae families of Caudovirales. Identification of auxiliary metabolic genes (AMGs) in sponge-associated viruses indicates that phages may complement the bacterial metabolic pathways [27]. Recently, a group of Ankyphages, encoding auxiliary ankyrin repeats (ANKs), was discovered. It was demonstrated *in vitro*, that ankyrin proteins from these phages reduce the phagocytosis of bacteria by macrophages, implicating that phages may mediate the symbiosis between bacteria and sponges [15]. Despite the several mechanisms discovered to date, the roles of viruses as a part of a sponge holobiont are not completely understood.

In this study, we aimed to characterize the interplay between microbial communities and viruses associated with three species of the White Sea sponges, namely *Isodictya palmata, Halichondria panicea*, and *Halichondria sitiens* (all belong to Demospongiae, subclass Heteroscleromorpha), and the surrounding seawater. Using the 16S metagenomics analysis we show that each species of the polar sponges is inhabited by highly diverse yet distinct microbial community. Using a shot-gun sequencing and assembly of metagenomes of the bacterial and viral fractions we analyzed both the diversity of viral contigs found in the metagenomes and the arsenal of bacterial defense systems. Our study shows that in contrast to previously studied tropical and temperate sponges, the abundance of defense systems appeared to be comparable for metagenomes of the White Sea sponges and surrounding water. Using the network-based approach for analysis of CRISPR-spacers found in CRISPR arrays and protospacers from viral genomes, we revealed possible phage bursts that originated in *I. palmata* and were further spread to *H. sitiens* and *H. panicea*. Our work provides the first insight into the metagenomics of the three cold-water sponges from the White Sea and paves the way for a comprehensive analysis of the interactions between microbial communities and associated viruses.

## 2. Materials and Methods

### 2.1. Sample collection

Two (for *Isodictya palmata*) or three (for *Halichondria sitiens* and *Halichondria panicea*) marine sponge individuals were collected by SCUBA divers at 5-7 m water depth at the N. Pertsov White Sea Biological Station (WSBS, 66.5527° N, 33.1033° E) in the Kandalaksha Bay of the White Sea (Russia) on August 10-20 of 2018. Samples were held separately in 5 L of sterile marine water (filtered through the 0.22 μm Sartorius filter) for 2 h at 5°C. Identification of sponge species was performed by zoologist Dr. Boris Osadchenko and further confirmed by 18S rRNA gene region amplification and sequencing. 3 L of surrounding seawater at the sampling site were collected in a sterile container and were immediately processed in the WSBS laboratory.

### 2.2. Bacterial and viral fractions isolation

For bacterial fraction isolation from sponges, a 1 cm^3^ fragment of sponge tissue was diced and fragmented by razor and forceps in 25 mL of sterile marine water. The obtained suspension was centrifuged at 200 g for 5 min. The supernatant was collected and centrifuged further at 3500 g for 10 min. The obtained pellet was defined as bacterial fraction and used for DNA isolation. Depletion of sponge cells and spicules in the bacterial fraction was confirmed by microscopy. The supernatant collected after the second round of centrifugation was further filtered using the 0.2 μm filter (Sartorius). Collected flow-throughs were pooled for replicates of the same sponge species and concentrated with Pellicon XL50 tangential filtering cassette with 100 kDa cutoff membrane (Millipore). The resultant concentrate (15-20 mL) was defined as a viral fraction.

To isolate bacterial fraction from marine water, 3 L of water were pre-filtered through the 5 μm filter (Sartorius). The flow-through was further filtered using the 0.2 μm filter (Sterivex). The fraction bound to the 0.2 μm membrane was defined as a bacterial fraction and was used for DNA isolation. The second flow-through was concentrated with the Pellicon XL50 cassette with 100 kDa cutoff (Millipore). The resultant concentrate (15-20 mL) was defined as viral fraction.

For collection of viral particles, PEG 8000 was added to 10% (w/v) and NaCl to 1 M to the viral fractions. Samples were incubated at 4°C for 1 h and centrifuged at 3500 g for 30 min. The supernatant was discarded, and the pellets were resuspended in 400 µL of STM buffer (100 mM NaCl, 10 mM MgSO4, 50 mM Tris-HCl pH 7.5). Samples were vortexed with 400 µL of chloroform and then centrifuged at 4°C for 5 min at 200 g. The resultant aqueous phase contained concentrated viral particles and was used for DNA isolation.

### 2.3. DNA extraction

DNA was extracted from bacterial fractions using the Diatom DNA Prep kit (Galart Diagnosticum) according to the manufacturer’s manual. For the bacterial fraction of marine water, settled in the 0.2 μm Sterivex filter, the filtering unit was opened, and the membrane was fragmented using the sterile razor. DNA was purified from the fragmented filter with the Diatom DNA prep kit. Then, DNA was extracted twice with phenol-chloroform and chloroform and precipitated in ethanol for the additional purification.

To purify DNA from the viral fractions, the samples (400 µL) were incubated with 1 µL of RNase A (Thermo Fisher Scientific) for 15 min at 37°C. Then, 50 µL of a lysis buffer was added (10% SDS, 20 mM EDTA, 200 mM Tris-HCl pH 7.5) and the samples were incubated with Proteinase K (Thermo Fisher Scientific), final concentration 100 µg/mL, for 1h at 55°C. DNA was extracted with an equal volume of phenol:chloroform:isoamyl alcohol (25:24:1, BioRad), and traces of phenol were removed by triple extraction with equal volumes of chloroform:isoamyl alcohol (24:1). Finally, DNA was precipitated with ethanol. DNA precipitates were dissolved in 20 µL of TE buffer and stored at −20°C. DNA concentration was assessed with Qubit (Invitrogen) and its integrity was checked by agarose electrophoresis.

### 2.4. High-throughput sequencing

For bacterial fractions from sponges and marine water, a V3-V4 region of 16S rRNA gene was amplified using standard degenerate primers fused with sequencing adapters (see Illumina guide for 16S Metagenomic Sequencing Library Preparation, Part # 15044223 Rev. B). The amplicon libraries were prepared and sequenced at Kurchatov Institute Core Sequencing Center using the 250+250 bp paired-end protocol with Illumina MiSeq.

DNA of bacterial or viral fraction replicates was pooled and submitted for shot-gun sequencing. Libraries were prepared at Skoltech Genomics Core Facility and were sequenced with 150+150 bp paired-end protocol using Illumina HiSeq.

### 2.5. 16S rRNA data analysis

Raw reads were trimmed and filtered using Trimmomatic version 0.39 (SE -phred 33 HEADCROP 17 ILLUMINACLIP:2:30:10 MINLEN:150) [28]. Survived forward reads were processed with DADA2 pipeline v. 3.6.2 [29] (including additional trimming, denoising, and errors correction) giving amplicon sequence variants (ASVs). The ASVs were clustered using MMseqs2 v. 10-6d92c [30] (coverage>0.95, identity >0.98) and representative sequences of clusters were further treated as operative taxonomic units (OTUs). OTUs were returned to DADA2 and taxonomy was assigned to OTUs using the SILVA SSU taxonomic training data formatted for DADA2 v.138 [31]. Finally, sequences classified as eukaryotes were removed. PCoA, alpha-diversity, and taxonomic analyses were performed with the phyloseq package v. 1.30.0 [32].

### 2.6. Shot-gun metagenomes assembly and annotation

Raw reads were quality checked with FastQC [33] and trimmed and filtered using Trimmomatic v. 0.39 (PE -phred 33 LEADING:3 TRAILING:3 ILLUMINACLIP:2:30:10 MINLEN:36). Survived read pairs and forward unpaired reads were assembled with SPAdes (with metaspades option on and k-mer length 55, 99, 121, 127) for *de novo* assembly [34]. The resultant assemblies were analyzed with QUAST [35]. Contigs longer than 5 kb were selected for further analyses. For taxonomy annotation, contigs were aligned against NCBI’s nr database using the DIAMOND blastx v. 0.9.24 [36]. The results were transferred to MEGAN v. 6.19.7 [37] and analyzed by the LCA algorithm. For accurate assembly of viral sequences, the survived reads were assembled with metaviralSPAdes tool [38]. In all selected bacterial and viral contigs ORFs were predicted by MetaGeneMark [39]. ORFs were annotated using DIAMOND blastp against NCBI’s nr database. Average coverage depth of contigs was obtained by mapping of reads used for *de novo* assembly on selected contigs with Bowtie 2 v. 2.4.4 [40], and computed from the bam files with SAMtools depth v. 1.10 [41].

### 2.7. Identification of viral sequences and taxonomic assignment

To detect viral sequences, contigs from metaviralSPAdes were processed using ViralVerify [38], VirSorter2 [42], and CheckV [43]. For ViralVerify detected sequences were analyzed with ViralComplete [38]. Sequences, recognized as viruses by at least two out of three pipelines were considered viral. The final set of sequences was clustered using Cd-hit v. 4.8.1 (coverage > 0.8, identity > 0.95) [44] to assemble a non-redundant set. Taxonomy of viral contigs was assigned by gene-sharing network approach using vConTACT2 v. 0.9.22 with default database Prokaryotic Viral RefSeq 207 and with ‘-s ‘MetaGeneMark’ --rel-mode ‘DIAMOND’ parameters [45]. The network was visualized using Cytoscape v. 3.8.0 [46]. Phylogenetic trees were constructed for viral contigs with the ViPTree web server [47].

### 2.8. Prediction and quantification of phage defense systems and AntiCRISPR proteins

Antiphage defense systems were detected in contigs using the PADS arsenal database [48]. Briefly, groups of orthologous sequences from PADS arsenal database were clustered using MMseqs2 with coverage>0.7, identity>0.85. Representative sequences were aligned using MAFFT with mafft-linsi (--maxiterate 100) [49] and obtained alignments were polished with trimAl (gap threshold 50%) [50]. HMM profiles were constructed from the polished alignments with HHMER v. 3.3.2 with hmmbuild algorithm [51]. ORFs from selected contigs were scanned with hmm profiles using hmmsearch and hits with e-value<10e-10 were selected. If an ORF was detected by several HMM profiles, a profile with the lowest e-value was considered. Putative defense systems were detected as at least two ORFs separated by less than 5 ORFs, which were recognized by two different HMM profiles belonging to one defense system in the PADS arsenal database. Additionally, defense systems were detected by PADLOC and DefenseFinder [52][53].

Raw counts of defense system genes and putative defense systems were obtained by simple counting of the genes and systems in the metagenomes.

For weighted and normalized quantification of putative defense systems or genes, an assembly length, total number of mapped reads, and coverage depth of contigs were considered, and the raw counts of defense system genes and putative defense systems were transformed according to the formula:

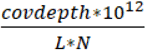, where L is the total length of a metagenome assembly, N is the total number of reads mapped to the assembly, covdepth is an average coverage depth of a contig, 10^12^ - is a scaling coefficient universal for all metagenomes used to obtain numerical value in a range of raw counts for defense genes and putative defense systems.

### 2.9. CRISPR spacer detection and analysis

CRISPR-Cas spacer arrays were detected with CRISPRCasTyper [54]. Retrieved sequences were clustered with Cd-hit (coverage > 0.95, identity > 0.95) to obtain a non-redundant dataset and to identify spacers shared between metagenomes. To identify spacers shared with CRISPRCasdb [55] they were clustered with sequences from the database using Cd-hit (coverage > 0.95, identity > 0.95). To detect possible protospacers, spacers were searched against the RefSeq viral sequences and viral assemblies derived from the White Sea metagenomes using BLASTN [56] with search parameters - word_size 8 -dust no -qcov_hsp_perc 95.

## 3. Results

### 3.1. Samples description and processing

Visually healthy individuals of marine sponges *Halichondria sitiens* (3 samples), *Halichondria panicea* (3 samples), and *Isodictya palmata* (2 samples), as well as one sample of surrounding seawater, were collected in August 2018 at the single site near N. Pertsov White Sea Biological Station (WSBS, **Figure 1A, 1B**). Bacterial fractions were extracted from the sponges and the seawater by step-wise centrifugation and filtration, respectively. Viral fractions were concentrated by tangential filtration from the flow-through after the bacterial fraction isolation. The composition of bacterial communities was investigated by amplification and sequencing (250+250 bp, Illumina MiSeq) of the V3-V4 region of 16S rRNA genes resulting in an average sequencing depth of 165 thousand reads per sample (**Figure 2D**). Sequencing data were processed by a combination of DADA2 pipeline with the subsequent analysis using the phyloseq R package (**Figure 1C**). Due to the inefficient merging of read pairs by DADA2, analysis was performed for forward reads only. To remove microvariations of ASVs and to work at a genus level, the ASVs were clustered to OTUs with a 98% identity threshold. The rarefaction curve analysis demonstrated that the communities were sequenced with a saturating sequencing depth (**Figure S1A**).

**Figure 1.**
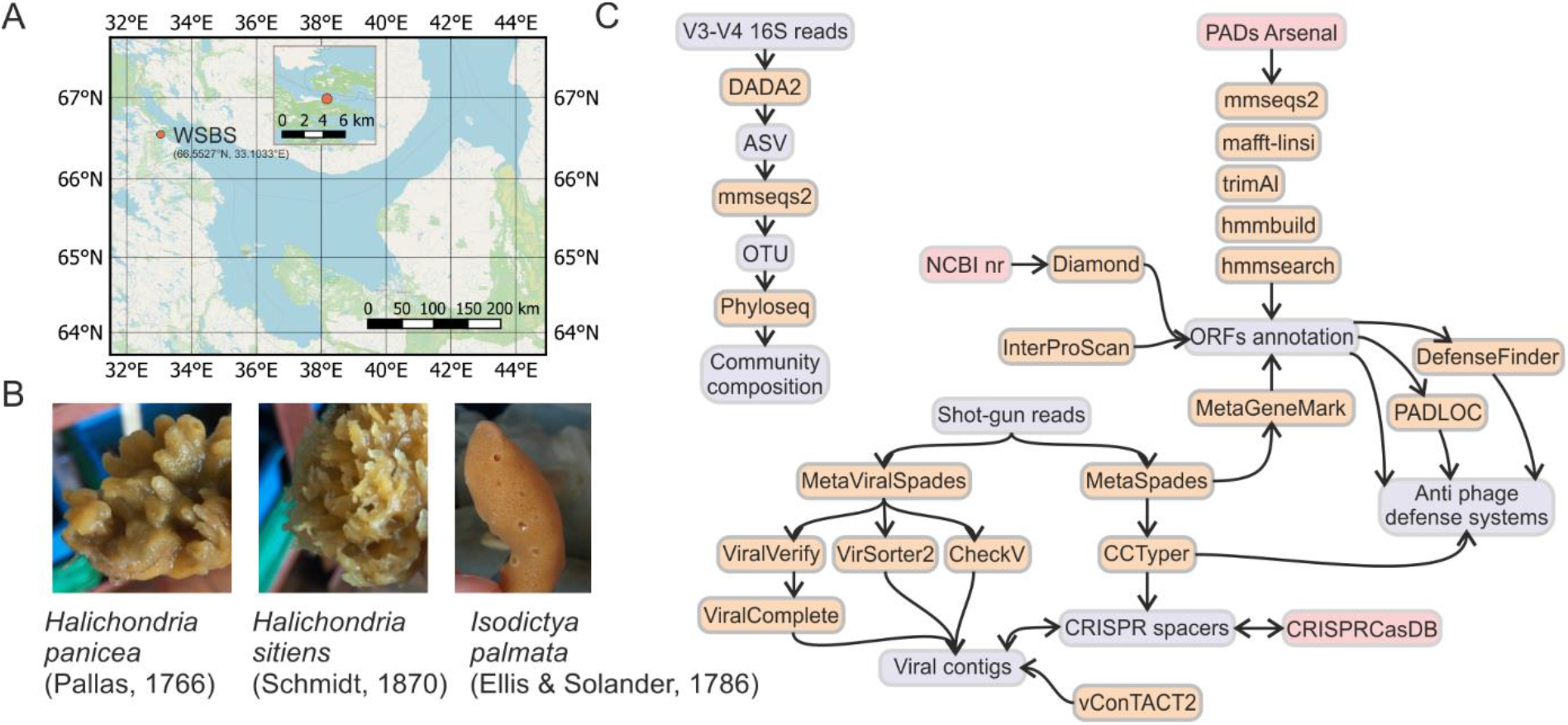
Samples collection and data analysis pipeline. (A) Samples of marine water and marine sponges were collected near the N. Pertsov White Sea Biological Station (WSBS), the Kandalaksha Bay of the White Sea, Russia. (B) Representative fragments of the collected sponges - *H. sitiens, H. panicea, I. palmata*. (C) Program pipelines used for data analysis.

**Figure 2.**
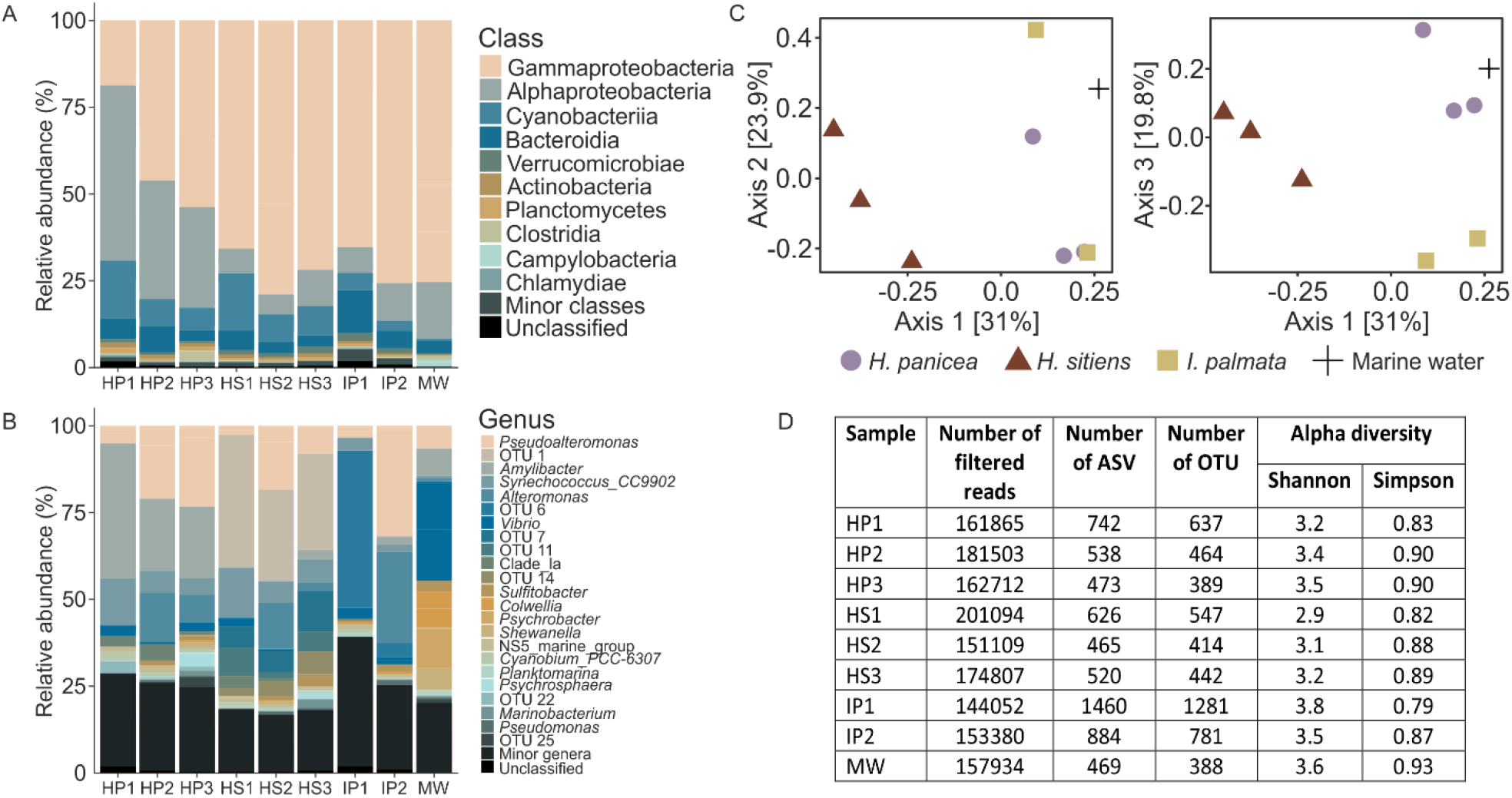
Diversity of White Sea sponge-associated and surrounding seawater microbial communities. (A) Taxonomic distribution of operational taxonomic units OTUs at the level of classes. *H. panicea* (3 replicates, HP), *H. sitiens* (3 replicates, HS), *I. palmata* (2 replicates, IP), marine water (MW). OTUs that were not assigned at the phylum level were grouped as Unclassified. (B) Taxonomic distribution of OTUs at the level of genera. OTUs that were not assigned at a genus level are indicated by numbers. The unclassified group as in A. (C) PCoA of Bray-Curtis dissimilarity between the studied microbial communities. (D) 16S sequencing statistics and alpha diversity metrics for the samples probed.

Total shot-gun metagenomes of the bacterial and viral fractions were sequenced using the Illumina HiSeq; on average, 120 mln high-quality reads were generated per metagenome (**Table 1**). Analysis of the cumulative length of metagenomes demonstrated the signs of length saturation, indicating the appropriate sequencing depth for the samples (**Figure S1B**). The overall data analysis pipeline, including assembly and annotation of metagenomes, identification of viral sequences and antiphage systems, and analysis of 16S data is shown in **Figure 1C**.

**Table 1.** Shot-gun metagenomes sequencing and assembly statistics. Bct and Vrs are for metagenomes of bacterial and viral fractions, respectively.

| Sample | Number of reads after filtering | Total number of contigs | Total assembly length, Mb | N50 | L50 | Number of contigs >5 kb | Total assembly length (contigs >5 kb), Mb |
| --- | --- | --- | --- | --- | --- | --- | --- |
| *H.panicea* (Bct) | 156744899 | 1048333 | 704.2 | 1678 | 62534 | 10147 | 102.6 |
| *H. sitiens* (Bct) | 151978145 | 660860 | 515.4 | 2309 | 36966 | 11252 | 105.4 |
| *I. palmata* (Bct) | 151705427 | 984734 | 626.6 | 1646 | 49855 | 8239 | 87.0 |
| Marine water (Bct) | 139611995 | 1409157 | 783.8 | 1271 | 74205 | 7 222 | 82.2 |
| *H.panicea* (Vrs) | 105797172 | 773423 | 477.2 | 1253 | 54605 | 3723 | 50.6 |
| *H. sitiens* (Vrs) | 81791648 | 416792 | 250.4 | 1313 | 21446 | 2579 | 33.8 |
| *I. palmata* (Vrs) | 74156030 | 214337 | 111.1 | 1227 | 8941 | 965 | 11.1 |
| Marine water (Vrs) | 107464178 | 1078397 | 691.0 | 1437 | 56832 | 7208 | 100.6 |

### 3.2. 16S metagenomics revealed distinct complex bacterial communities associated with the White Sea sponges and marine water

Metagenomic analysis of the V3-V4 fragment of 16S rRNA gene revealed the difference in the compositions of communities associated with marine sponges and marine water. The most abundant phyla in sponges were Proteobacteria (80%), Cyanobacteria (9%), and Bacteroidetes (6%) (**Figure 2A**). The most abundant classes in *H. panicea* were Gamma- and Alphaproteobacteria (40% and 38%, respectively), followed by Cyanobacteria (10%). *H. sitiens* and *I. palmata* were largely dominated by Gammaproteobacteria (72% and 71%, respectively), Alphaproteobacteria (8% and 9%), and Cyanobacteria (11% and 4%). On a genus level, *Pseudoalteromonas* OTUs were present in all samples, while sponges were dominated by sponge-specific OTUs not abundant in marine water (**Figure 2B**). For *H. panicea* the characteristic OTU was OTU 2 classified as *Amylibacter* (27%), for *H. sitiens* it was OTU 1 (31%, order UBA10353), and for *I. palmata* it was OTU 6 (25%, unclassified Gammaproteobacteria). The *Amylibacter* OTU had a 100% identity with *Candidatus Halichondribacter symbioticus*, a recently described symbiont of *H. panicea* from Icelandic waters [57]. Correspondently, the results of the PCoA analysis using Bray-Curtis dissimilarity demonstrated that sponges are inhabited by specific communities differ from surrounding marine water (**Figure 2C**). On average, 497 OTUs were detected in *H. sitiens*, 468 in *H. panicea*, 1031 in *I. palmata*, and 388 in marine water (**Figure 2D**). Alpha-diversity Shannon and Simpson indexes indicated that all bacterial communities of sponges and sea water had a complex structure, but they were dominated by specific groups of microorganisms (**Figure 2D**).

### 3.3. Enrichment and diversity of viromes from the White Sea

To obtain viral particles from the sponge and marine water samples, an enrichment strategy was developed, which is based on the fractionation of samples by consecutive centrifugation steps and/or filtering (see Materials and Methods). Shot-gun metagenomes of the resultant viral fractions were sequenced and assembled (**Table 1**), and viral sequences were identified by several algorithms including ViralVerify, VirSorter2, and CheckV. To reduce the number of false positives, sequences detected by at least two pipelines were considered as putative viral sequences and used in further analysis (**Figure S3A**). Applying this approach, in total 453 viral sequences were identified in metagenomes. An average number of retrieved viral sequences was considerably higher for the viral fractions than for the bacterial fractions (96 vs. 27, t-test p-value 0.027), indicating an efficient enrichment procedure (**Figure 3A**). Interestingly, the number of viral sequences shared between viral and bacterial metagenomes was low (6 sequences, **Figure 3B**, Venn diagram) allowing us to suppose that the enrichment method could be selective for some viral particles. Viromes associated with particular sponge species and marine water were distinct, as indicated by the low number of viral sequences shared among samples (**Figure 3B**, heatmap). This observation is not surprising given the distinctive profiles of bacterial communities revealed in these samples using 16S metagenomics (**Figure 2B-C**).

**Figure 3.**
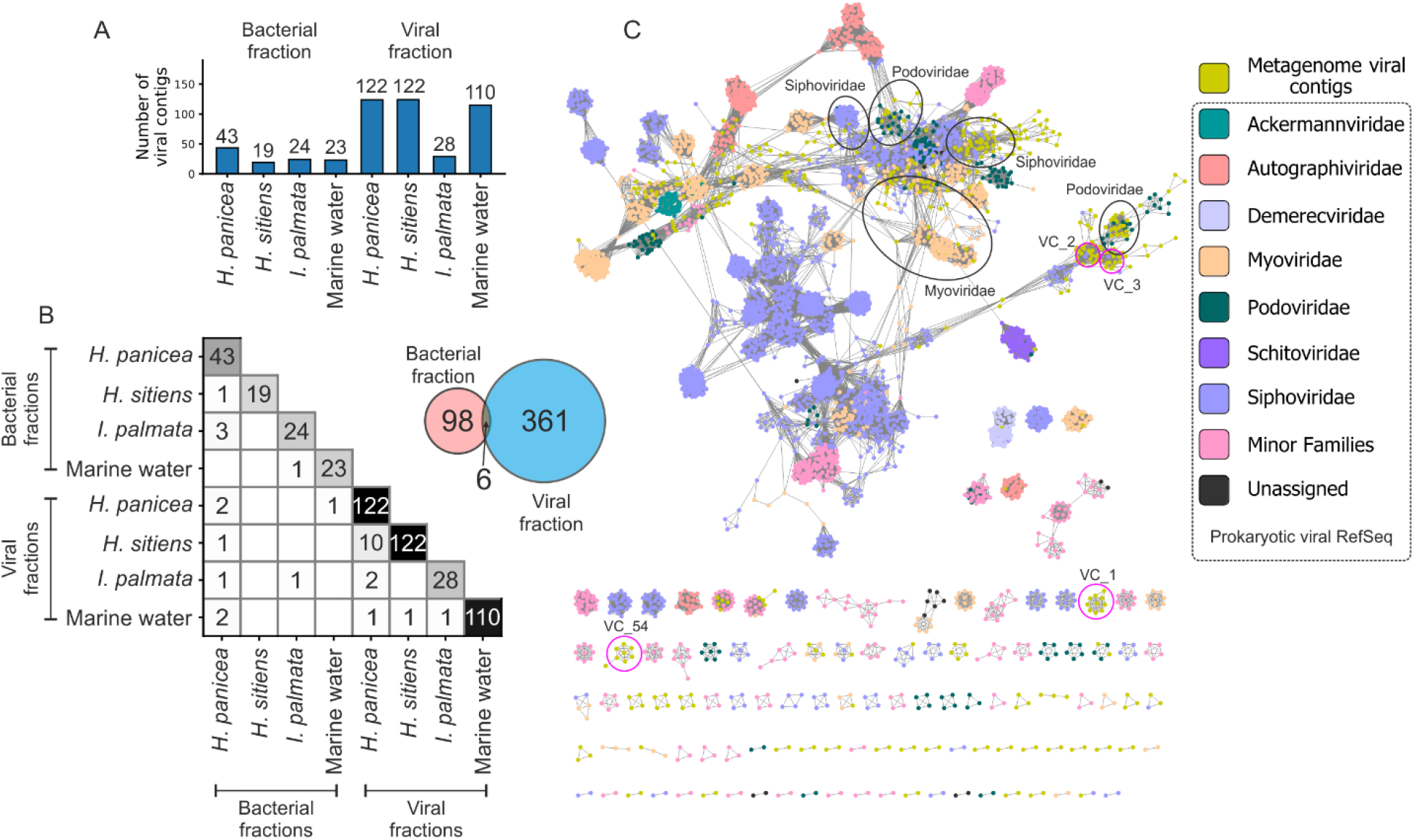
Analysis of viromes associated with marine sponges and sea water. (**A**) The number of viral contigs identified in different samples. Viral contigs were clustered using CD-hit (identity >95%, coverage >80%). (**B**) Viral contigs shared between the samples. The number of viral contigs shared between bacterial and viral fractions. (**C**) Sequence similarity network (SSN) constructed by vConTACT2 with Viral RefSeq-prokaryotes database clustered with viral contigs detected in the White Sea metagenomes. Nodes and edges represent the viral genomes and a significant similarity shared between them, respectively. Black ovals indicate viral contigs derived from the metagenomes in clusters with phages from the families of Caudovirales. Pink circles mark the most abundant viral clusters composed exclusively of viral contigs (VC) from the White Sea metagenomes. The network was visualized in Cytoscape.

Phylogeny of retrieved viruses was investigated using vConTACT2 (**Figure 3C**). From 453 identified viral contigs, only 10% (46) were clustered to the sequences from the reference database. These belong to the order Caudovirales and families Myoviridae (21 sequences), Siphoviridae (19 sequences), and Podoviridae (6 sequences). 90% of viral sequences had no similarity to known viral sequences; among them, 261 sequences were clustered (81 clusters) and others were assigned as outliers or singletons (**Figure S3B**).

### 3.4. Antiphage systems detected in metagenomes

Putative defense genes and defense systems were detected in assembled and annotated metagenomes using the custom HMM profiles. The profiles were built from the alignments constructed using the sequences from the PADS Arsenal database [48]. By simple counting, the most abundant putative defense genes belonged to SEPTU, BREX, ZORYA, and DISARM systems (**Figure 4A**). The most abundant putative defense systems included RM, ZORYA, and CRISPR-Cas (**Figure 4B**). Normalization of metagenomes and adjustment of gene copy-number by the average coverage depth of contigs did not change the abundance of putative defense genes and systems dramatically, except that ABI replaced CRISPR-Cas in top-three defense systems (**Figure 4C-D**). In accordance with the custom search results, the most prevalent systems identified in metagenomes by DefenseFinder were RM, followed by ABI and CRISPR-Cas (**Figure S4A**). Analysis with PADLOC also demonstrated the high prevalence of CRISPR-Cas systems in the metagenomes (**Figure S4B**).

**Figure 4.**
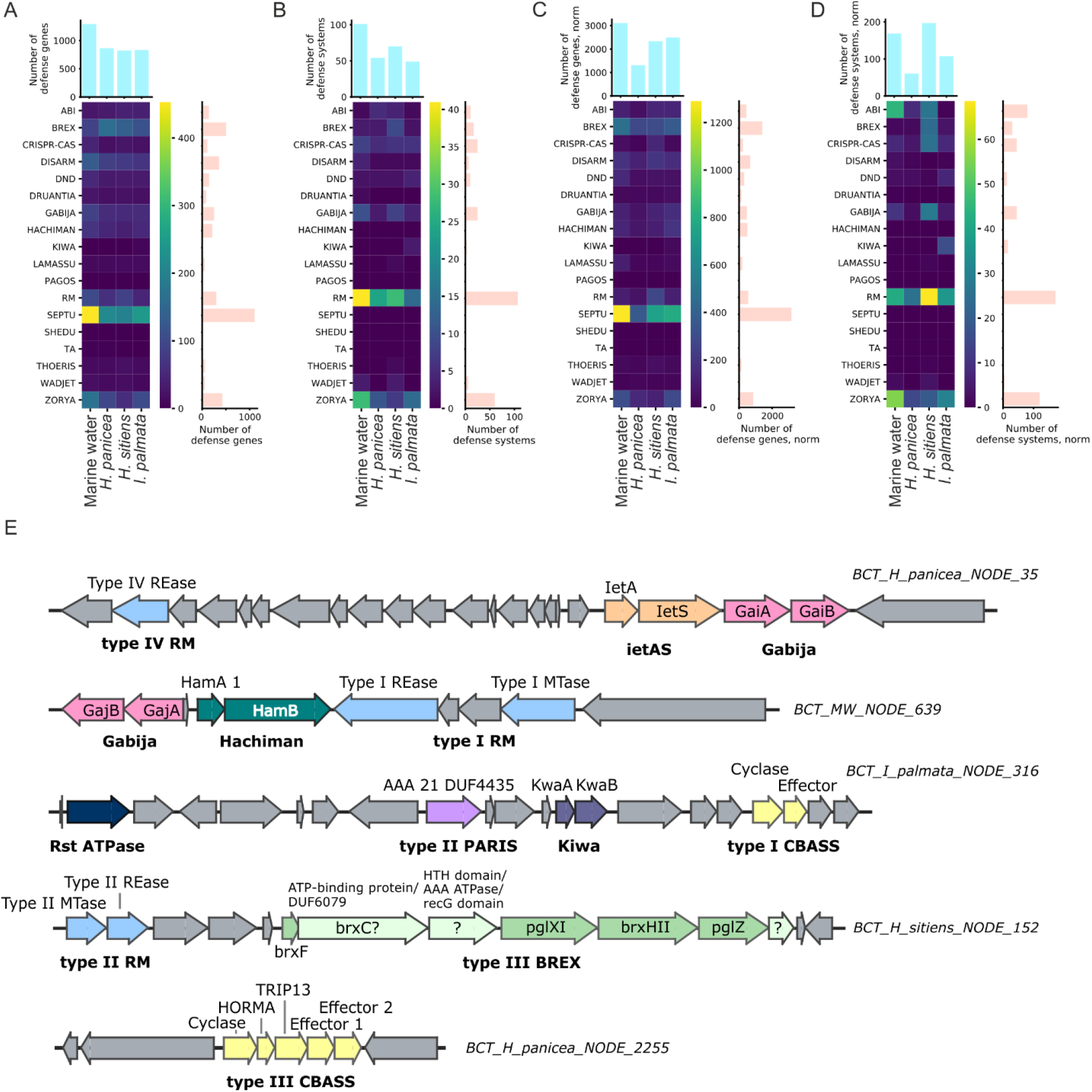
Heatmaps representing the abundance of putative defense system genes (A) and putative defense systems (B) found in metagenomes. Heatmaps representing the normalized abundance of putative defense system genes (C) and putative defense systems (D) found in metagenomes. (E) Examples of genomic islands enriched in defense systems genes and unusual defense systems identified in the metagenomes of White Sea sponges and marine water.

Overall, among the studied metagenomes, marine water had slightly increased raw counts of defense systems and is particularly rich in RM, ZORYA, Gabija, and CRISPR-Cas systems (**Figure 4B, S4A**). PADLOC also identified a substantial number of the CBASS systems in the marine water metagenome (**Figure S4B)**. After the normalization procedure, however, the metagenome from *H. sitiens* was found to have the highest number of defense systems (**Figure 4D**), thus indicating the presence of abundant bacteria enriched with the RM, CRISPR-Cas, Gabija, Gao, and QatABCD defense systems.

In the metagenomes from *H. panicea* and the marine water, two unusual type-III CBASS systems were found. These systems contain two effector genes with a limited similarity shared, a feature previously not described for the known CBASS systems [58] (**Figure 4E**). Also, for several contigs, most of which were assigned to Gammaproteobacteria by MEGAN, islands including co-clustered defense systems were identified. In one of the defense islands from the *H. sitiens* metagenome, a novel variation of the type-III BREX system was found. It contains a non-canonical *brxC* gene with limited homology to the classical C-genes and the insertion of an additional gene of unknown function between the *brxC* and *brxX* genes (**Figure 4E**).

### 3.5. Analysis of CRISPR spacers show the flow of viruses between marine communities

CRISPR immune systems are known to keep the record of invasions of bacteria by phages and other mobile elements in the associated CRISPR arrays [59]. Non-array sequences matching the spacers from arrays, known as protospacers, evidence the facts of invasion and the subsequent immune adaptation of the host.

We used CRISPRCasTyper to detect CRISPR arrays in metagenomes and derived spacers. Noteworthy, the metagenomes of viral fractions contained microbial DNA. Totally, 2358 unique spacers were identified in 197 CRISPR arrays, of which 171 were found in the bacterial fractions, and 26 in the viral fractions. None of them matched spacers from the CRISPRCasDB database (contains 291402 unique spacers) (**Figure 5C**). The spacer sets from different White Sea metagenomes poorly overlap, thus highlighting the distinct compositions of bacterial communities (**Figure 5A**). Interestingly, metagenomes of viral fractions had spacers not detected in bacterial fractions, which may originate from arrays of small-size bacteria passing through the 0.22 μm filter (**Figure 5B**).

**Figure 5.**
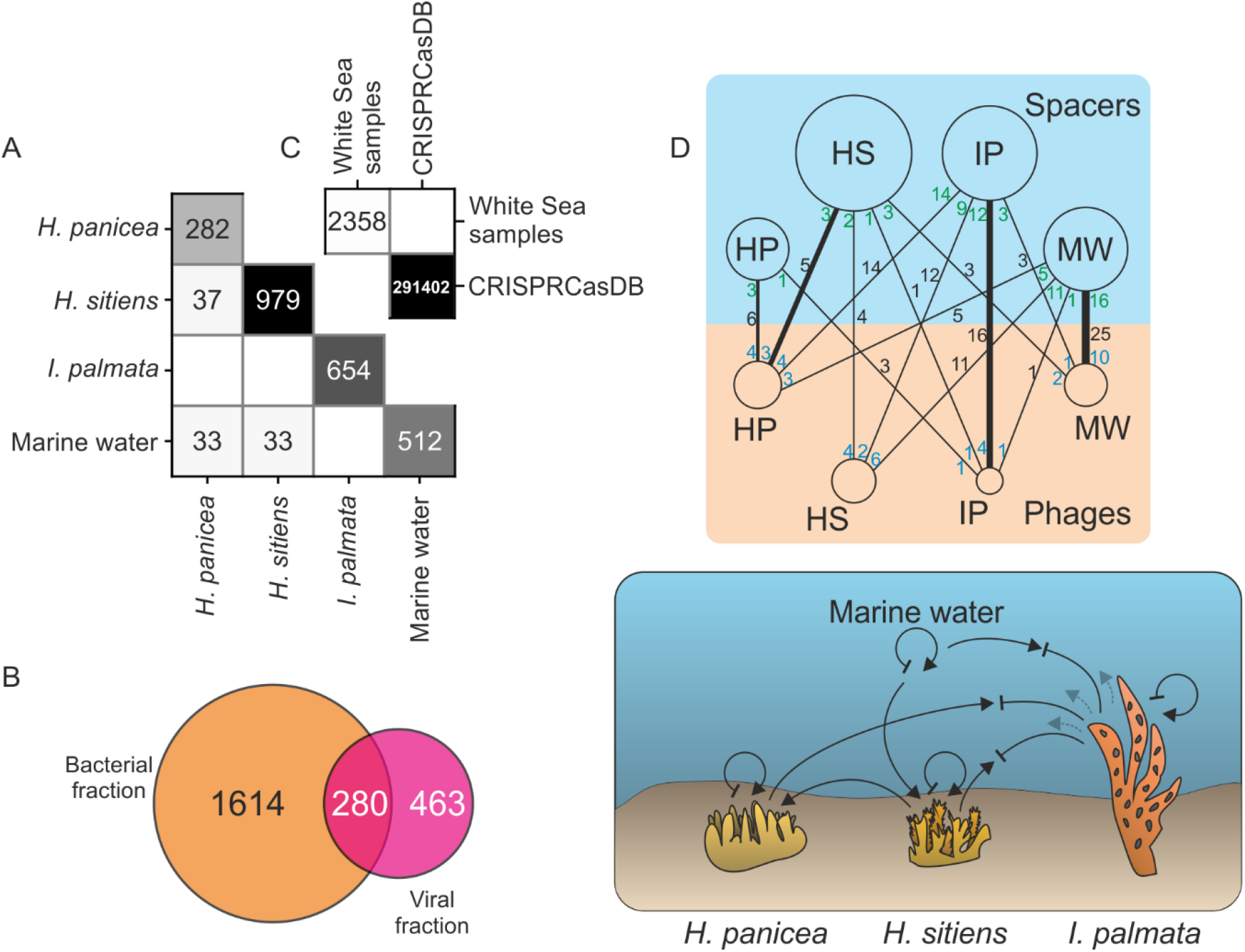
(A) Numbers of CRISPR spacers shared between studied bacterial communities. (B) The number of CRISPR spacers shared between bacterial and viral fractions. (C) The number of CRISPR spacers identified in the White Sea metagenomes shared with CRISPRCasDB database. (D) Top, a bipartite community-community network of virus-host interactions. The upper and the lower nodes represent the source of the spacers and the viral contigs, respectively. The area of nodes is proportional to the number of spacers or viral contigs identified in metagenomes. Black, green and blue numbers on the edges of the network account for the total number of matches between spacers and protospacers, number of matching spacers, and number of viral contigs with protospacers, respectively. For each of the upper nodes, an edge with the largest number of spacers is shown in bold. Arrows represent a phage burst or a phage flow between communities; inhibition arrows indicate the defense against phages. Dark black and pale arrows correspond to more recent and older events, respectively.

To detect potential protospacers, we searched spacer sequences in the RefSeq Viruses database and the set of putative viral contigs. Only one reliable match was found with a genome of Pseudomonas phage phi2 from the RefSeq Viruses, and 121 matches were found with putative viral contigs discussed above. Next, we analyzed the sources of spacers and protospacers and constructed a bipartite community-community network of virus-host interactions (**Figure 5D**). As can be seen from the network, the number of productive interactions (captured by the spacer-acquisition events; black numbers on the edges of the network) were typically higher within communities than between different communities. As an exception, the spacers from the bacterial community of *H. sitiens* more frequently matched with viral contigs from *H. panicea* than with any other community, including the *H. sitiens* itself, potentially indicating the flow of viruses between the communities. Generally, *H. panicea* contained more viruses matched by spacers from other samples than any other community, possibly reflecting the ongoing spread of viruses. In contrast, *I. palmata* had more spacers matching viruses from other samples than any other community, likely highlighting the recent numerous events of a successful defense. Supporting this speculation, *I. palmata* metagenomes showed the smallest number of putative viral contigs (**Figure 3A**). Additionally, the average coverage depth of a ubiquitous 144-145 kb-long viral contig, closely related to *Pseudoalteromonas* phages from the Myoviridae group (**Figure S5A**), was considerably lower in the *I. palmata* metagenome (9 times less than in *H. panicea* and 36 times less than in *H. sitiens*). The drop in the contig abundance can be explained by efficient CRISPR adaptation and interference, as 10 different spacers, matching this contig (the CRISPR system I-F type, **Figure S5B**), were found in *I. palmata* metagenome (**Figure S5C**). Of note, 9 out of 10 protospacers had an adjacent CC PAM classical for the I-F systems [60]. Taken together, our data indicate that a directional flow of viruses may exist between closely localized bacterial communities.

## 4. Discussion

To investigate possible interactions between bacteria and viruses within and between sympatric marine sponges, bacterial and viral fractions from three species of White Sea sponges and surrounding marine water were isolated and their metagenomic DNA was sequenced.

### 4.1. Sponge species specificity of associated bacterial and viral communities

Microbiomes of *H. panicea* from different geographical locations were investigated earlier by strain culturing [61], 16S metagenomics [7,62], and whole metagenome sequencing [63]. In accordance with the previously published 16S metagenomics data, Alphaproteobacteria, in particular, a few sponge-specific OTUs from this class (e.g., *Amylibacter*), were prevalent in *H. panicea*-associated bacterial communities from the White Sea. The *Amylibacter* OTU sequence was identical to the 16S of the symbiotic bacterium *Candidatus Halichondribacter symbioticus*, recently described in *H. panicea* from Icelandic waters [57], indicating the wide geographical association between the sponge and the symbiotic bacterium. It is worth noting that Gammaproteobacteria were unusually prevalent in the White Sea *H. panicea* samples (especially the genera *Pseudoalteromonas, Alteromonas*, and *Vibrio*). We speculate that this reflects the community profile shift due to the increased water temperature during the summer season of 2018 [64].

In the current study, the compositions of microbial communities of *H. sitiens* and *I. palmata* were investigated using metagenomic techniques for the first time. The communities were largely dominated by Gammaproteobacteria, Alphaproteobacteria, and Cyanobacteria, a pattern, observed for many sponge species [10,65]. At the genus level, communities were dominated by specific yet poorly taxonomically assigned OTUs (OTU 1 for *H. sitiens* and OTU 6 for *I. palmata*), which arguably correspond to novel putative sponge symbionts. The predominance of a single bacterium has been shown for several different sponges previously [57,66,67].

Only a minor fraction of viral contigs identified in the White Sea metagenomes were classified to known groups of viruses. These groups were mostly represented by Myoviridae, Podoviridae, and Siphoviridae, the phage families typically found in viromes of different sponges from various habitats [15,26,27,68]. The majority of viral contigs did not fall into clusters with known reference sequences, representing the viral “dark matter” [69].

The compositions of microbial metagenomes, viromes and spaceromes (the set of all CRISPR spacers) were compared between different sponge species. We revealed a prominent host species specificity in all cases. This observation highlights the distinct nature and independence of sponge-associated communities, even spatially closely localized [6,70].

### 4.2. Abundance of the defense systems, unusual CBASS and BREX systems

Sponges filter thousands of liters of water a day, accumulating virus particles that can affect the composition of the associated bacterial communities [71]. Thus, sponge-associated microorganisms are forced to provide reliable protection against phages and plasmids in conditions of high viral load and high cell density [16,17,19,21,25,72]. To investigate the defense potential of the communities, we performed mining of the White Sea sponge and marine water metagenomes and revealed a variety of defense systems. Unexpectedly, the diversity and abundance of the systems in the sponge and marine water metagenomes were comparable. This observation argues with the published data that postulate the higher prevalence of various defense systems in the metagenomes of sponge-associated communities over the surrounding water (19 of 20 sponge metagenomes studied). Noteworthy, this rule was also correct for the two species of cold-water Antarctic sponges [20]. However, in the sponge *Scopalina ruetzleri*, collected in the equatorial Atlantic, CRISPR-Cas and RM systems were less abundant, indicating possible deviations from the general scheme [19]. One can speculate that the discrepancy can be explained by the skew of the earlier studies toward the detection and counting of CRISPR-Cas, RM, and TA systems. We hypothesize that the defense potential of the bacterial communities of cold-water sponges could be compensated by diverse defense systems belonging to other and/or yet unknown types. A unified analysis, including the detection of an extended set of defense systems, is needed for the correct comparison of the metagenomes.

Cyclic-oligonucleotide-based anti-phage signaling systems (CBASS) are a group of bacterial antiphage systems [73]. Upon phage infection, the CBASS generates signaling cyclic oligonucleotides. In turn, the oligonucleotides activate an effector, which promotes cell death resulting in abortive infection. The effectors can contain various cell-killing domains: patatin-like phospholipase, endonuclease, peptidase, etc. [73]. We identified unusual type-III CBASS variants in two contigs from marine water and *H. panicea* sponge: clusters contained two effector genes, both encoding putative DNA endonucleases. Interestingly, the effectors were distant homologs, which probably implies a different substrate specificity or, alternatively, the effectors can be activated by different signaling oligonucleotides.

Bacteriophage exclusion defense system (BREX) blocks phage replication and discriminates between host and phage DNA by methylation patterns [74,75]. A novel variation of the type-III BREX cluster with an unknown gene inserted between *brxC* (putative ATPase) and *brxX* (putative methyltransferase) was identified. Using HHpred we predicted that this protein contains a helix-turn-helix domain on the N-terminus and its C-terminal part is homologous to ATP-dependent helicases. Also, it contains a conserved triad typical for the Mg^2+^-binding site, which is a common characteristic for some ribonucleases [76,77]. Recently, insertion of a type-IV RM nuclease named *brxU* was observed in a plasmid-borne type-I BREX system between the *brxC* and *brxX* genes [78]. We speculate that there could be a hotspot for the insertions of accessory proteins, such as nucleases, inside the BREX cluster. The hotspot may be mediated by the frequent rearrangements occurring at the *brxX* gene that might be due to the high toxicity of its product [74].

### 4.3. Tracking the viral flow between communities by a spacer-protospacer network

Bacteria acquire new spacers when a CRISPR-Cas adaptation complex captures fragments of a phage genome and incorporates them into CRISPR arrays during the process known as adaptation. When the phage infects the cells harboring the matching spacer, the viral genome is recognized and degraded by a CRISPR interference complex guided by the spacer transcript [79,80]. The information stored in CRISPR-Cas arrays can be used to reveal the history of the phage-host interactions [81]. Based on this, we investigated the exchange of viruses between sponge-associated communities. Using a network approach, we identified several examples of inter-community viral exchanges. In particular, in the *I. palmata* metagenome, a contig containing the type-IF CRISPR-Cas system and an associated array was found to carry spacers matching the abundant viral contigs identified in all investigated sponges. Although these viral contigs were highly prevalent in the case of *H. panicea* and *H. sitiens*, no matching spacers were found in the microbial metagenomes of these organisms. In contrast, the viral contigs were considerably less abundant in *I. palmata*, implying the effective interference by the CRISPR-Cas system. We speculate that this phage could have been first propagated in *I. palmata*, where it was suppressed by the activity of CRISPR-Cas. At the same time, it was spread and invaded the communities of other nearby sponges not yet adapted to it. Interestingly, the I-F array contained two groups of spacers targeting the phage genome - at the beginning and the end of the array. The first group contained more spacers perfectly matching the genome. This observation and the fact that new spacers are usually incorporated at the beginning of an array [82] suggest that the community of *I. palmata* faced the phage twice - earlier and, probably, during the current phage outburst. Our network-based approach can be scaled up and applied for the investigation of viral exchanges within large sets of bacterial communities.

## 5. Conclusions

This study presents the first insight into the bacterial and viral communities associated with three cold-water marine sponge species from the White Sea. We demonstrated that despite the close spatial proximity of the holobionts on the seafloor, the sponge-associated communities are species-specific and differ substantially by a composition of bacterial OTUs, diversity of associated viruses, and acquired CRISPR-spacers. At the same time, the frequent exchange of viruses between the communities exists, as indicated by the acquisition of CRISPR-spacers against viruses from another community. The abundances and diversity of defense systems in the metagenomes of sponges and the metagenome of water are comparable, which could be a specific feature of polar ecosystems. Finally, in the metagenomes, we identified unusual variations of BREX and CBASS bacterial defense systems. With this study, we begin to fill the gaps in the knowledge of complex interactions between bacterial and viral communities associated with polar marine organisms.

## Supplementary Materials

### Funding

The work was supported by a grant from the Ministry of Science and Higher Education of Russian Federation (agreement Nº075-10-2021-114 from 11.10.2021). D.S. was supported by the Russian Foundation for Basic Research grant (project No. 20-34-90069). Sequencing at Skoltech Genomics Core Facility was supported by the Skoltech Life Sciences Program grant.

## Acknowledgments

We thank the personnel of the White Sea Biological Station, particularly, Dr. Alexander Tzetlin and Dr. Tatyana Neretina for the opportunity to collect and process samples at WSBS, WSBS SCUBA diving team for samples collection, and Dr. Boris Osadchenko for identification of sponge species.

## Author Contributions

A.R. processed samples, analyzed data, and drafted the manuscript. V.F. collected and processed samples. S.T. prepared the 16S rRNA sequencing libraries and sequenced the samples. S.D. acquired funding and finalized the manuscript. D.S. conceptualized and designed the research, analyzed data, prepared figures, and drafted the manuscript. All authors read and approved the final manuscript.

## Institutional Review Board Statement

Not applicable.

## Informed Consent Statement

Not applicable.

## Data Availability Statement

Sequencing data were deposited in SRA (*ID_to_be_assigned*). Metagenome assemblies were deposited in GeneBank under the BioProject *ID_to_be_assigned*. Custom code used for data analysis is available from https://github.com/sutormin94/WSBS_Sponges_metagenomics.

