## Supplementary Figures for "An interplay between viruses and bacteria associated with marine sponges from the White Sea revealed by metagenomics"

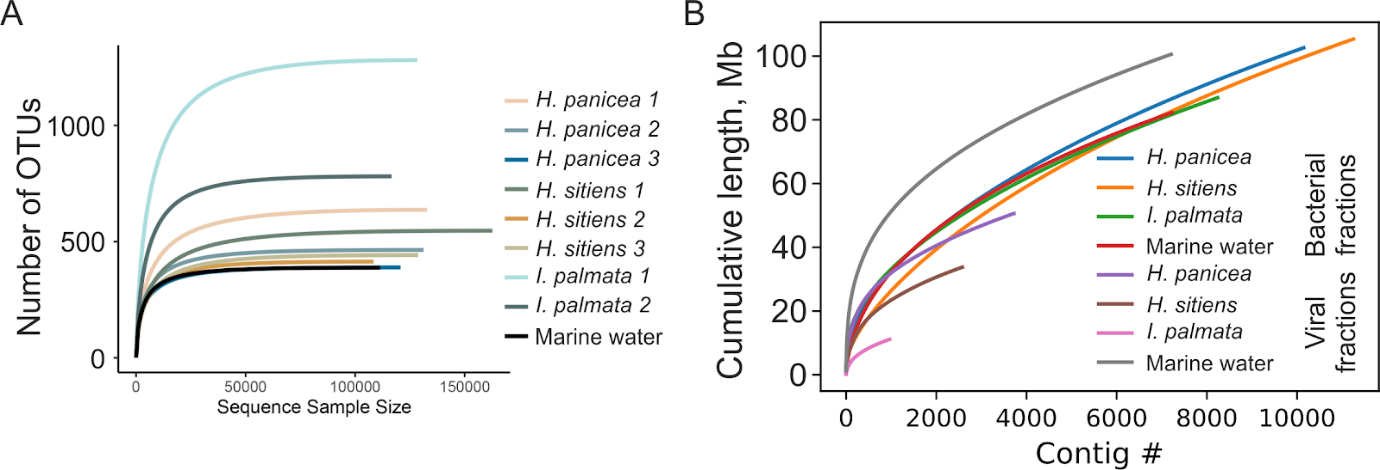

**Figure S1**. (**A**) Rarefaction curves analysis of V3-V4 16S metagenomes sequences in this study. (**B**) Analysis of the cumulative length of metagenomes for contigs > 5 kb.

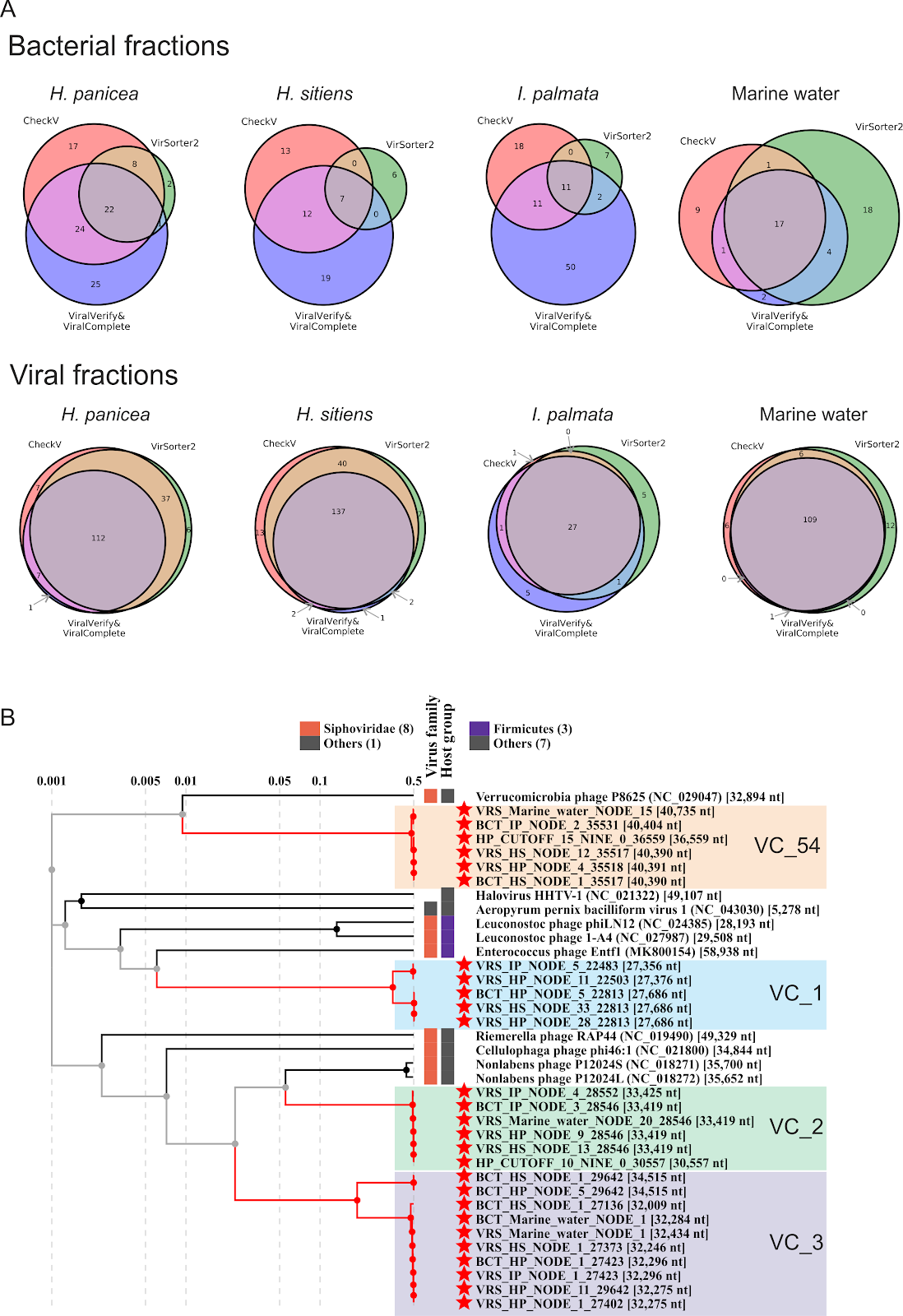

**Figure S3**. (**A**) The Venn diagrams represent the concordance between the predictions of viral sequences by ViralVerify & ViralComplete pipeline, VirSorter2, and CheckV. Numbers are given for sequences before clustering with Cd-hit. Sequences detected by at least two tools were defined as putative viral contigs and selected for further analyses. (**B**) Phylogenetic tree of representative viral contigs (indicated by red asterisks) belonging to 4 the most abundant viral clusters composed exclusively of viral contigs from the White Sea metagenomes. The tree was prepared using ViPTree. Viral cluster’s IDs correspond to the IDs indicated on the network in Figure 3C.

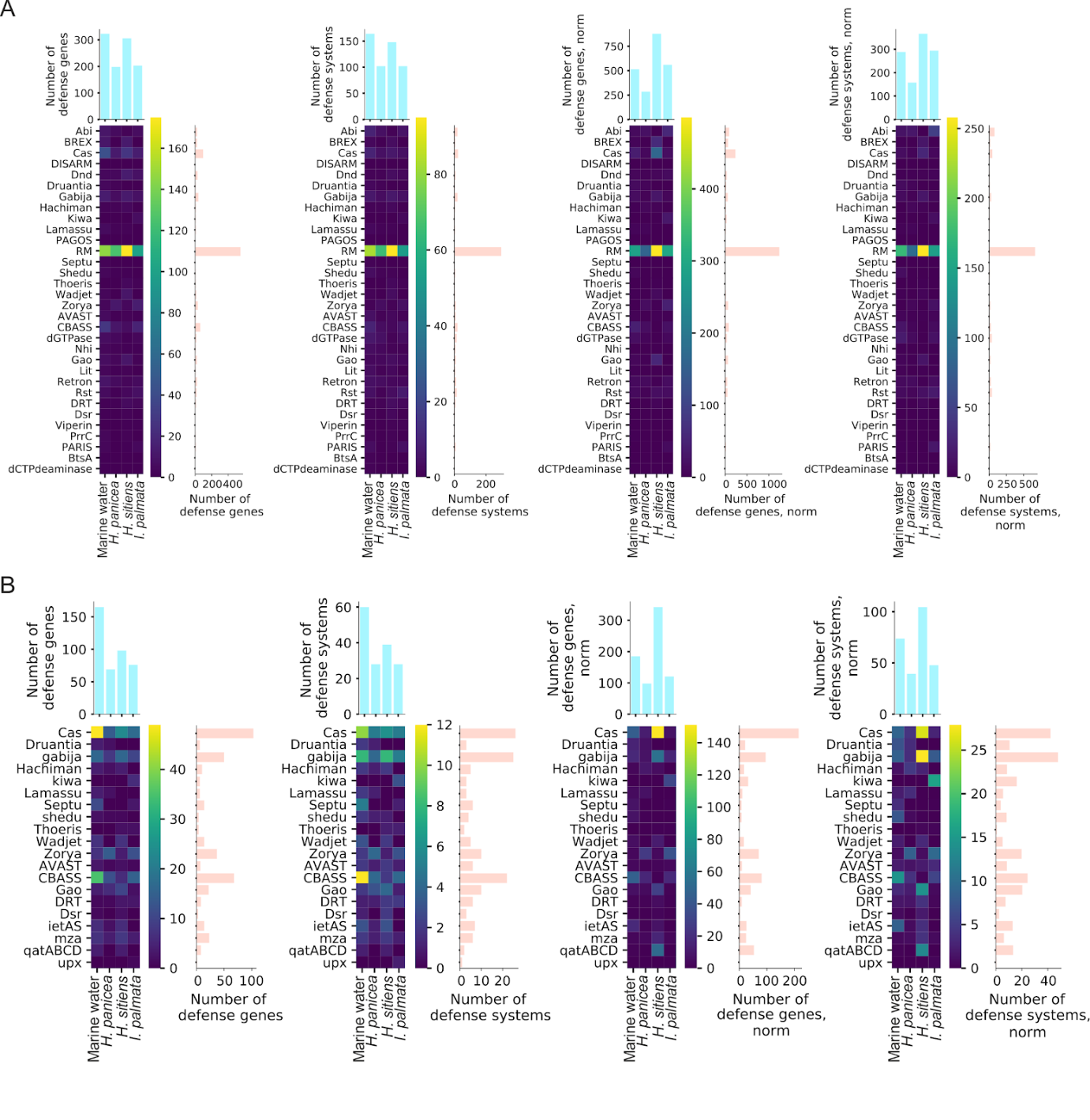

**Figure S4**. (**A**) Heatmaps representing, from left to right, the numbers of defense systems genes, numbers of putative defense systems, normalized numbers of defense systems genes, and normalized numbers of putative defense systems found in metagenomes using DefenseFinder. (**B**) Heatmaps representing, from left to right, the numbers of defense systems genes, numbers of putative defense systems, normalized numbers of defense systems genes, and normalized numbers of putative defense systems found in metagenomes using PADLOC.

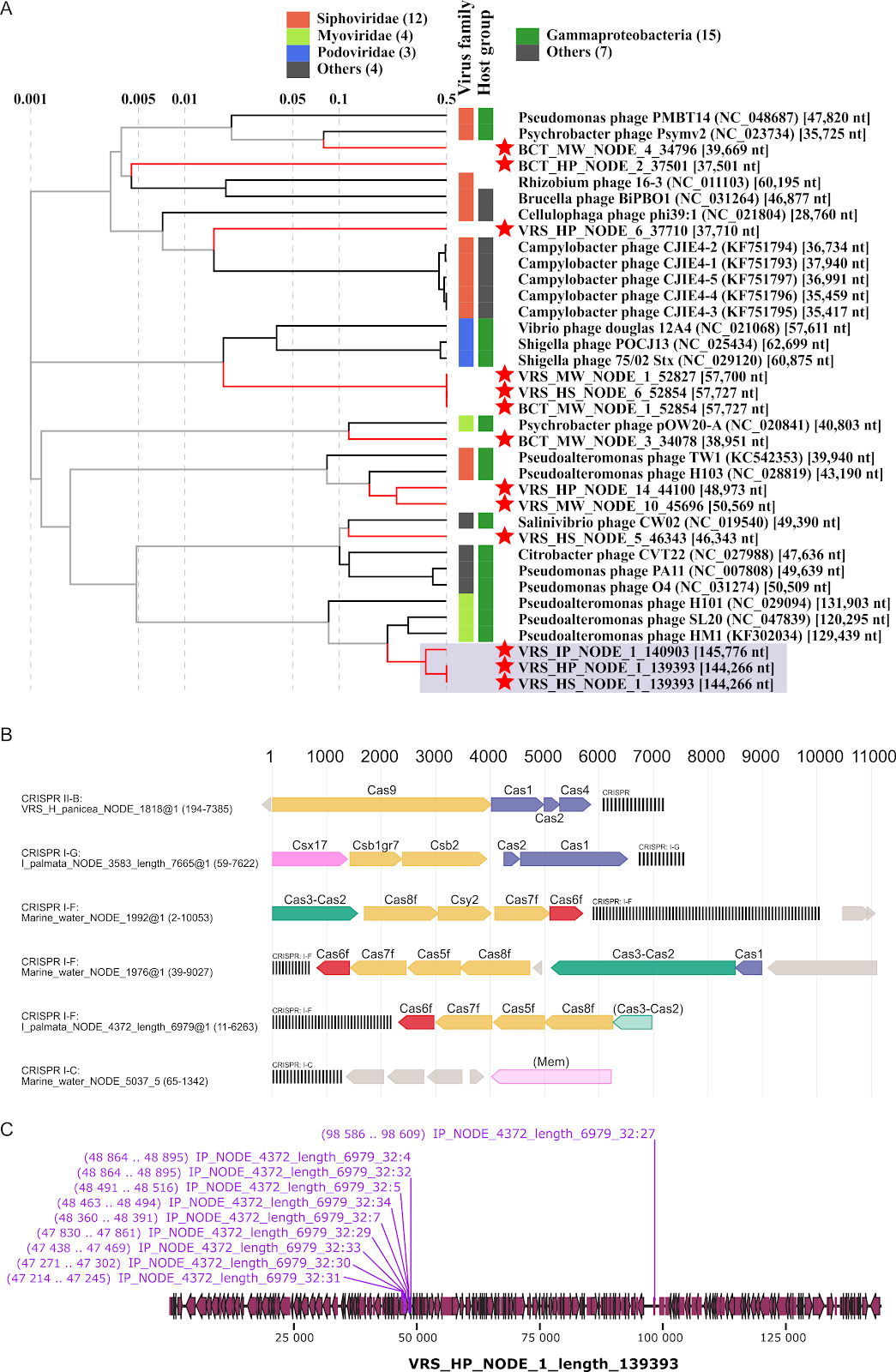

**Figure S5**. (**A**) Phylogenetic tree of viral sequences constructed with ViPTree. Putative viral contigs with detected protospacers are highlighted by red asterisks. The blue rectangle marks the group of related contigs found in all sponges, containing protospacers matching by several spacers of the I-F CRISPR-Cas system from the *I. palmata* metagenome. (**B**) CRISPR-Cas systems containing spacers for which matching protospacers were found in the putative viral contigs. (**С**) Annotation of a viral contig targeted by 10 spacers (positions of protospacers are labeled) from I-F CRISPR-Cas system identified in the *I. palmata* metagenome.
